# The Functional RNA Identification (FRID) Pipeline: Identification of Potential Pseudoknot-Containing RNA Elements as Therapeutic Targets for SARS-CoV-2

**DOI:** 10.1101/2023.04.03.535424

**Authors:** Peter C. Forstmeier, McCauley O. Meyer, Philip C. Bevilacqua

## Abstract

The COVID-19 pandemic persists despite the development of effective vaccines. As such, it remains crucial to identify new targets for antiviral therapies. The causative virus of COVID-19, SARS-CoV-2, is a positive-sense RNA virus with RNA structures that could serve as therapeutic targets. One such RNA with established function is the frameshift stimulatory element (FSE), which promotes programmed ribosomal frameshifting. To accelerate identification of additional functional RNA elements, we introduce a novel computational approach termed the Functional RNA Identification (FRID) pipeline. The guiding principle of our pipeline, which uses established component programs as well as customized component programs, is that functional RNA elements have conserved secondary and pseudoknot structures that facilitate function. To assess the presence and conservation of putative functional RNA elements in SARS-CoV-2, we compared over 6,000 SARS-CoV-2 genomic isolates. We identified 22 functional RNA elements from the SARS-CoV-2 genome, 14 of which have conserved pseudoknots and serve as potential targets for small molecule or antisense oligonucleotide therapeutics. The FRID pipeline is general and can be applied to identify pseudoknotted RNAs for targeted therapeutics in genomes or transcriptomes from any virus or organism.

## Introduction

As of March 2023, the COVID-19 pandemic has led to over 675 million infections and 6.9 million deaths worldwide (Dong et al.) and continues to pose a threat despite the development of novel mRNA vaccines (Jackson et al. 2020; Krienke et al. 2021; Muik et al. 2022). Several small molecules that target disease-related RNAs have recently been advanced (Ingemarsdotter et al. 2018; Costales et al. 2020; Haniff et al. 2020; Zeller et al. 2022). However, while there have been advances for antivirals that target the SARS-CoV-2 proteins such as the RNA-dependent RNA polymerase (RdRp) (e.g. remdesivir, molnupiravir) and the main protease (M^pro^ or nsp5) (e.g. paxlovid and nirmatrelvir-ritonavir) (Simonis et al. 2021; Marzi et al. 2022; Saravolatz et al. 2022), there are not antivirals that directly target the RNA genome of SARS-CoV-2 (Wu et al. 2020; Simonis et al. 2021; Marzi et al. 2022; Saravolatz et al. 2022). Indeed, the lifecycle of SARS-CoV-2 is regulated by the formation and stability of diverse RNA structures (Manfredonia et al. 2020; Cao et al. 2021; Huston et al. 2021; Sun et al. 2021; Lan et al. 2022), which can perform essential biological activities that mediate gene regulation (Winkler et al. 2004; Peselis and Serganov 2014).

Functional RNA elements are typically characterized by strong secondary structure and the presence of one or more pseudoknots—structural elements in which the nucleobases contained within a hairpin loop, internal loop, or large bulge form non-nested Watson-Crick or wobble base pairs with nucleobases outside of the hairpin (Figures 1 and S1) (Staple and Butcher 2005). Due to this non-nested “reaching” topology, pseudoknots compact an RNA’s structure, creating sites for catalysis and for binding of small molecules (Herschlag 1998). Indeed, riboswitches and ribozymes almost always have one or more pseudoknot(s) (Staple and Butcher 2005; Peselis and Serganov 2014). Additionally, proteins can bind to certain pseudoknot-containing RNAs to effect frameshifting (Namy et al. 2006; Giedroc and Cornish 2009; Ketteler 2012).

**Figure 1:**
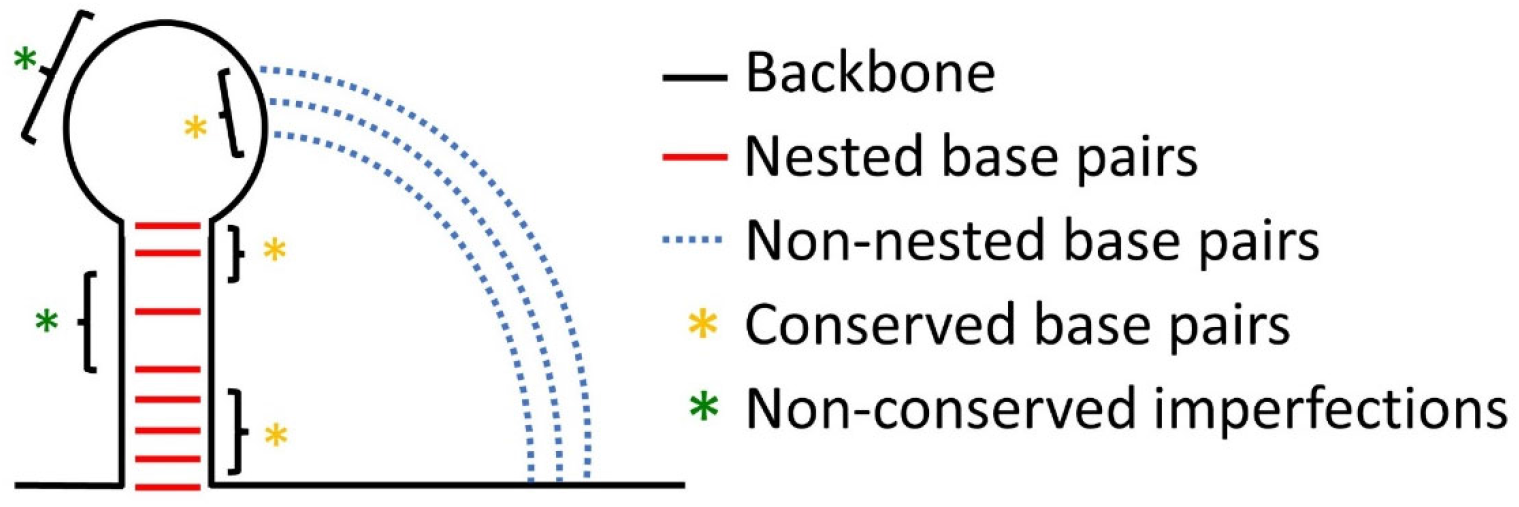
Depiction of a simplified functional RNA element. The backbone is black, nested base pairs are red, and non-nested base pairs are blue dotted lines. Features are marked with potential regions of higher (yellow) and lower (green) conservation. In this example, the apical and basal regions of the stem are more conserved (yellow).

Proper function of an RNA is intimately linked with proper folding of its structure (Doherty and Doudna 2000; Hofacker et al. 2006; Kobori and Yokobayashi 2016; Meyer et al. 2020), which creates a selective pressure to conserve the structure, including its pseudoknots (Figure 1). Oftentimes, base identity is less important than base pair conservation (Shang et al. 2012). Because functional RNA elements are structured and compact, they are often good targets for small molecule therapeutics, which can interrupt a viral lifecycle and prevent replication of the virus (Hargrove 2020). Small molecules can lock an RNA into an inactive conformation or recruit factors to degrade the RNA (Kaul et al. 2004; Costales et al. 2018; Crooke et al. 2018; Meyer et al. 2020). In addition, antisense oligonucleotides (ASOs) can target specific RNAs via sequence complementarity, interrupting the base pairing structure of the functional RNA element to inhibit its activity (Bennett and Swayze 2010; Khvorova 2022; Zhu et al. 2022). Pseudoknots are particularly attractive targets for therapeutics (Zafferani and Hargrove 2021). On the one hand, pseudoknots can be locked into a single folded conformation, preventing the flexibility needed for functions like frameshifting. For instance, Park et al. identified small molecules that bind specifically to the FSE of SARS-CoV and reduce its plasticity using *in silico* docking (Park et al. 2011; Ritchie et al. 2014). On the other hand, pseudoknots could be forced into the unfolded open hairpin state and rendered nonfunctional by the binding of small molecules and antisense oligonucleotides into hairpin loops, internal loops, and bulges that could block pairing of the non-nested strand. Indeed, formation of non-native, alternative structures by the hairpin portion of a pseudoknot often limits the formation of the pseudoknot, underscoring the basis for such a pseudoknot displacement strategy (Pan and Woodson 1998; Chadalavada et al. 2002).

Given that RNA sequences adopt distinct conformations, each of which with a unique folding free energy, single nucleotide polymorphisms (SNPs) can shift the populations of various states (Halvorsen et al. 2010). Those SNPs that cause changes to the secondary structure of an RNA are referred to as “riboSNitches” (Ritz et al. 2012) and can have profound impacts on disease; for instance, they have been implicated in hyperferritinemia cataract syndrome, β-thalassemia, and hypertension (Halvorsen et al. 2010). Recently, we found that SNPs can act as riboSNitches in plants with some SNPs favoring their adaptation to local climates (Ferrero-Serrano et al. 2022). In this study, we introduce the term “pseudoSNitch” for those SNPs that change the structure of a pseudoknot. Indeed, riboSNitches and pseudoSNitches may have pronounced effects in RNA viruses such as SARS-CoV-2, which depend on RNA structures to regulate their lifecycle.

Using the plethora of available genomic data on SARS-CoV-2, we identify potential novel functional pseudoknot-containing elements. To accomplish this, we built the Functional RNA Identification (FRID) pipeline, leveraging existing programs and writing new ones as needed. The FRID pipeline’s predicted structure of the FSE was similar to its canonical structure (Zhang et al. 2021), which serves as an important positive control; additionally, we identified several naturally occurring, low-frequency mutations in the FSE that may influence its conformation and stability. The FRID pipeline also identified 8 non-conserved and 14 conserved pseudoknot-containing RNA elements, which may serve as therapeutic targets. Remarkably, approximately half of the FRID-predicted functional RNA elements align well with available *in vivo* experimental RNA structure probing data supporting their folds. The functional RNA elements identified herein serve as potential targets for development of small molecule and ASO therapeutics.

## Results and Discussion

The SARS-CoV-2 genome is ∼30 kb in length (Andrews et al. 2021). Its first open reading frame, ORF1a, encodes for a long polyprotein that contains the multidomain protein nsp3 (Lei et al. 2018; Lv et al. 2022; Puray-Chavez et al. 2022), while the genome’s last quarter encodes a variety of structural proteins that arise from subgenomic fragments containing a common 5′-end termed the transcription regulatory sequence leader (TRS-L) (Figure 2) (Alexandersen et al. 2020; Kim et al. 2020). To regulate production of non-structural proteins and the ratio of the ORF1a and ORF1ab gene products, the SARS-CoV-2 genome contains an AU-rich ‘slippery sequence’ and a frameshifting element (FSE) (Figure 2). The FSE is a well-characterized functional RNA with several stem-loops and a pseudoknot that together with the slippery sequence promotes programmed (–1) ribosomal frameshifting to facilitate translation of the downstream portion of the ORF1ab polyproteins.

**Figure 2:**
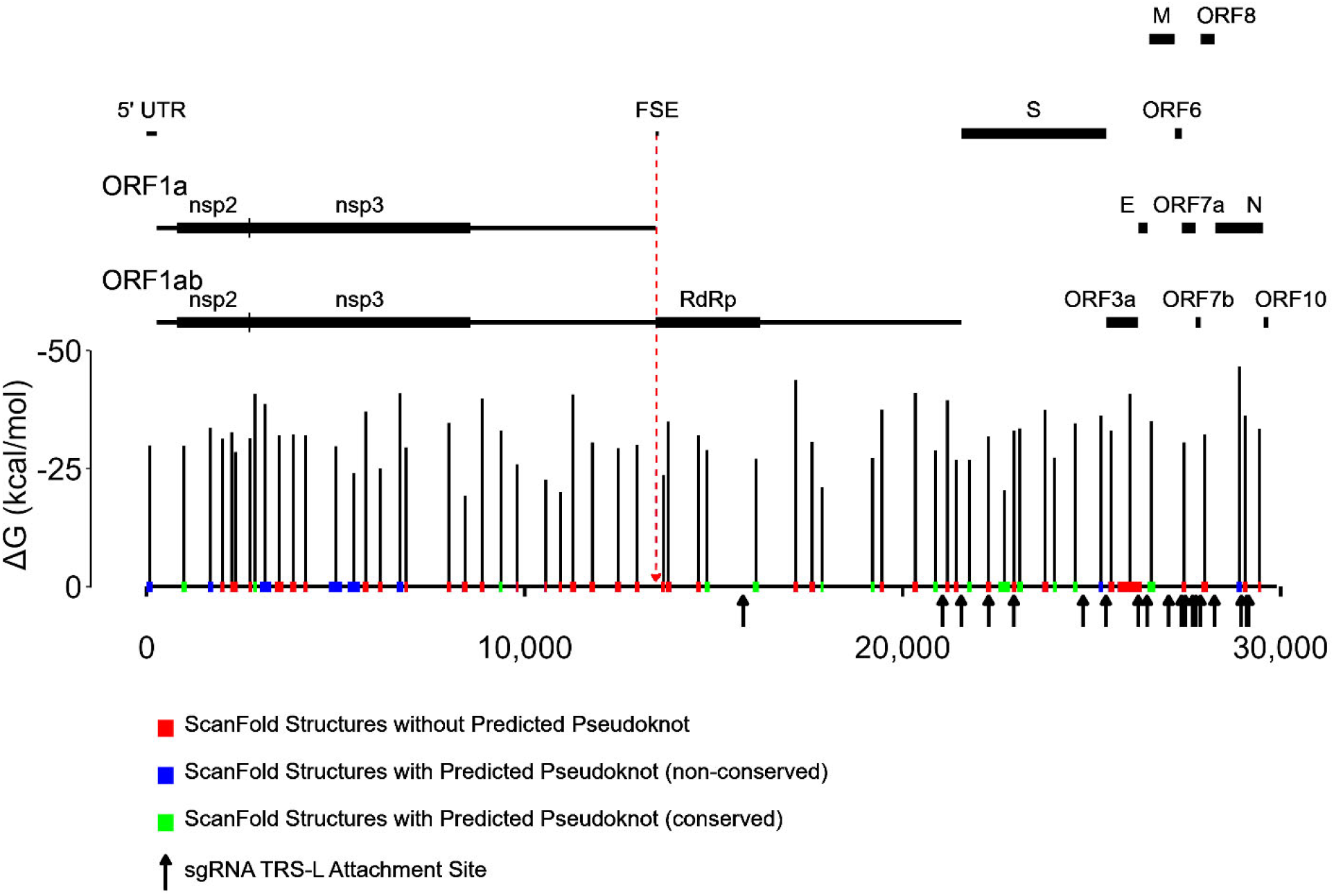
Output of the FRID pipeline mapped to the SARS-CoV-2 genome. The ScanFold-predicted structured regions without pseudoknots are red (N=38), with non-conserved pseudoknots are blue (N=8), and with conserved pseudoknots are green (N=14). The coloring and positioning of structures are the same as in Table S1. Each structured region has a vertical black line indicating its Mean Free Energy (MFE) ΔG value (in kcal mol^-1^). The vertical red dotted line indicates the location of the FSE. Arrows indicate the 3ʹ TRS-L junction sites of the top 25 most common sgRNAs (Kim et al. 2020). The following features are annotated: frameshifting element (FSE), spike protein (S), envelope protein (E), matrix protein (M), nucleocapsid protein (N), and open reading frames (ORFs). Only select nsp’s are depicted here.

To identify novel functional RNA elements in the SARS-CoV-2 RNA genome, we developed the Functional RNA Identification Pipeline (FRID). Our goal was to locate high probability functional RNA candidates for therapeutic targeting. The FRID pipeline was assembled in two modules, the Pseudoknot Module (PK Module) and the RNA Structure Conservation Module (RSC Module) (Figure 3). The PK Module predicts pseudoknot-containing RNA structures in the genome, while the RSC Module uses the output of the PK Module to assess the conservation of primary, secondary, and pseudoknot structure. We begin with sections on the development and optimization of these two modules. We then provide sections on application of the FRID pipeline, one focusing on the FSE and the other on identifying novel functional RNA elements within the SARS-CoV-2 RNA genome.

**Figure 3:**
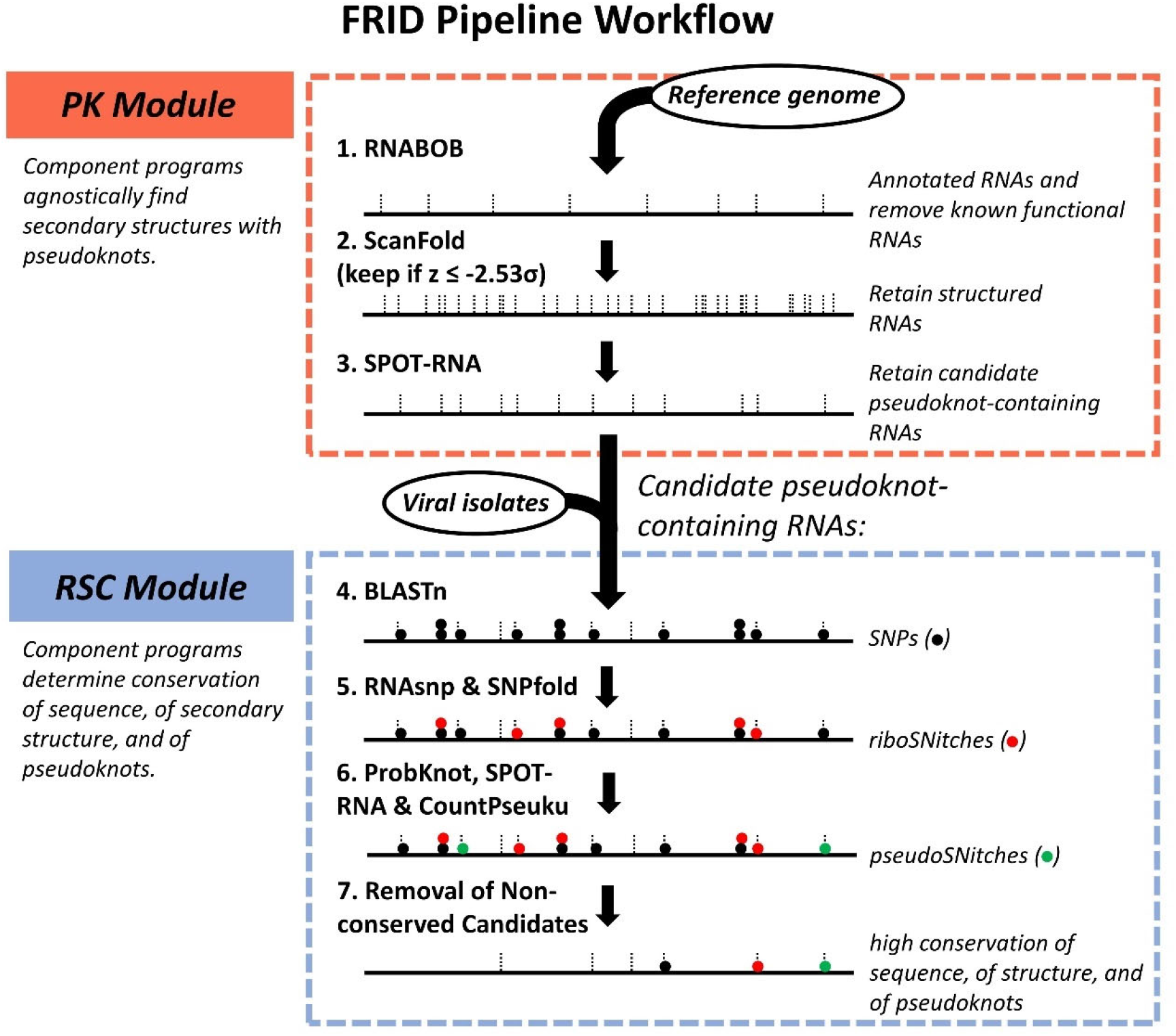
Overview of the Modules of the Functional RNA Identification pipeline (FRID). The pipeline is comprised of (1) the Pseudoknot (PK) Module (orange) and (2) the RNA Structure Conservation (RSC) Module (blue). The outputs of the component programs of each module are annotated in italic font to the right of each program. The small vertical dotted lines denote local structures. SNPs are represented by black circles, riboSNitches by red circles, and pseudoSNitches by green circles. The lines, SNPs, riboSNitches, and pseudoSNitches are for illustrative purposes.

### Development and Optimization of the Pseudoknot (PK) Module

The PK Module (Figure 3, orange dashed box) identifies pseudoknot-containing RNA elements and consists of three component programs: RNABOB (Eddy 2012), ScanFold (Andrews et al. 2018), and SPOT-RNA (Singh et al. 2019). Here, it identifies pseudoknot-containing elements in the reference genome of SARS-CoV-2 (NC_045512.2) by scanning it for and setting aside known functional RNAs using RNABOB (Figure 3, Step 1), scanning it for and retaining unknown structured RNAs using ScanFold (Figure 3, Step 2), and scanning those for the subset of pseudoknot-containing RNAs using SPOT-RNA (Figure 3, Step 3).

The first component program, RNABOB (Eddy 2012) (Figure 3, Step 1), scanned the SARS-CoV-2 RNA genome for known classes of riboswitches and ribozymes using RNA motif descriptors (see Supplemental Descriptors). We wrote custom descriptors for the deoxyguanosine, adenine, and guanine riboswitches, as well as the HDV, hammerhead, and twister ribozymes (Kingsford et al. 2007; Eddy 2012). One could, of course, include descriptors for a much wider range of RNA motifs, such as hairpins, ribozymes, and riboswitches (Bevilacqua and Blose 2008; Campillos et al. 2010; Mccown et al. 2017; Kavita and Breaker 2022). After validating our RNABOB descriptors on a set of positive and negative control RNAs, we scanned the SARS-CoV-2 genome for these motifs. Although we did not identify any of them, this screen is generally useful for identifying known classes of functional RNAs that will be assigned and set aside.

The second component program, ScanFold (Figure 3, Step 2) (Andrews et al. 2018), works without specifying an RNA motif descriptor. Instead, it searches the RNA genome agnostically for strong RNA structures, assigning each a z-score (Eq. S1) as a measure of the likelihood of a structure being non-random and therefore functional. We applied ScanFold to the SARS-CoV-2 RNA genome to identify those RNA elements with a significant z-score, defined herein as having a z-score of -2.53σ or lower (see Supplemental Text Sections 1 and 2, Figure S2, Tables S1-3, Eq. S2). We identified 60 such structured regions, which were mapped to the RNA genome (Figure 2), with their sequences and positions provided in Table S3.

As seen in Figure 2, these structured regions are dispersed throughout the genome but cluster primarily in its first and last quarters. The first quarter of the genome codes for the nsp3 multifunctional protein, whose production might be regulated by strong RNA structure, while the last quarter codes for various structural proteins, including the spike (S) protein, which are produced via discontinuous transcription of subgenomic (sg) RNAs (Kim et al. 2020).

Intriguingly, in the last quarter of the genome we predicted regions of strong RNA structure that cluster near or overlap with the TRS-L points of attachment to the sgRNAs (Figure 2, arrows) (Alexandersen et al. 2020). The mechanism for attachment of the TRS-L has been proposed to involve pausing of negative sense RNA synthesis (Kim et al. 2020). Perhaps the strong positive sense RNA structures predicted herein, which serve as the template for negative sense RNA synthesis, play a role in such pausing.

We briefly consider the consequences of adjusting the z-score threshold to a less stringent value. Lowering the z-score threshold, on the one hand, would generate more true positive candidates, but it would also generate many more false positives (Table S2). For instance, by using a threshold of just -2.10σ (instead of -2.53σ), the likelihood of finding a true positive becomes 100% in our practice set, which represents an increase in the number of true positives from 18 at z ≤ -2.53σ to 23 at z ≤ -2.10σ. However, this occurs at the expense of the likelihood of a predicted positive being a false positive increasing, which represents an increase in the number of false positives from 409 at z ≤ -2.53σ to 752 at z ≤ -2.10σ. In other words, 343 more false positives are obtained to get just five more true positives. Nonetheless, in some instances a less stringent threshold may be desirable; for instance, by lowering the threshold stringency to z ≤ -2.29σ, we were able to identify the FSE (see *Analysis of the Frameshifting Element (FSE) in SARS-CoV-2 Genome* below). Additionally, high throughput experimental techniques, such as genome-wide structure mapping (Rouskin et al. 2013; Ding et al. 2014) or massively parallel oligonucleotide synthesis (Carlile et al. 2019) could be used to screen thousands of candidates at once. Finally, it is worth considering the positive predictive value (PPV), which is the likelihood of a prediction being correct (Table S2). Not surprisingly, the PPV is maximal at the more stringent thresholds; in this example at a threshold of -2.92σ (Table S2). Thus, seeking more true positives by using a less stringent threshold comes at the expense of lowering the PPV.

Known functional RNAs typically contain one or more pseudoknots (Staple and Butcher 2005); therefore, we employed a pseudoknot searching component program, SPOT-RNA (Figure 3, Step 3) (Singh et al. 2019). This program was chosen because it had the highest PPV (i.e. was best at determining whether a positive is a true positive, albeit by a small margin) and was the most sensitive (i.e. was best at determining the most true positives here by a wide margin) of the three pseudoknot-predicting programs we tested (See Supplemental Text Section 3). However, SPOT-RNA is computationally expensive; we therefore searched only the 60 regions identified by ScanFold (Figure 2 and Table S3). We allowed G•U wobbles to count as pseudoknot base pairs because we found that these mutations retain functionality in saturation mutagenesis studies on ribozymes (See Supplemental Text Section 3, Figure S3, and Tables S4-5) (Kobori and Yokobayashi 2016). We found that 22 of the 60 structured regions have putative pseudoknots. These 22 candidate functional RNAs cluster in the coding regions of the nsp3 multifunctional and S proteins and are also present in the 5ʹ-UTR and RdRp elements (Figure 2, Table S3). The 22 RNA elements with predicted pseudoknots were then input into the RSC Module.

### Development and Optimization of the RNA Structure Conservation (RSC) Module

The RSC module (Figure 3, blue dashed box) determines which of the above-identified candidate pseudoknot-containing RNA elements identified in the PK Module (22 here) are strongly conserved and therefore potential candidate functional RNAs. To do this, more than 6,000 viral isolates were input into the RSC module and their conservation of primary sequence, secondary structure, and pseudoknot structure(s) was monitored via six component programs. In a region of interest, we judged conservation of sequence by the number of SNPs (Figure 3, Step 4), of secondary structure by the number of SNPs that were riboSNitches (Figure 3, Step 5), and of pseudoknot structure by the number of SNPs that were pseudoSNitches (Figure 3, Step 6). As a benchmark, we first calculated the level of conservation of a known functional RNA element in SARS-CoV-2, the FSE. Subsequently, candidate structured RNA elements with conservation levels equal to or higher than the FSE were considered putatively functional (Figure 3, Step 7).

The first three steps of the RSC module identify SNPs and classify them as riboSNitches, pseudoSNitches, or neither (Figure 3, Steps 4-6). We identified SNPs by comparing the viral isolates to the reference sequence using BLASTn (Zhang et al. 2000; Morgulis et al. 2008) (Figure 3, Step 4), and more than 50,000 SNPs were identified (provided in FRID_SNPs_archive.csv). To assure that only quality sequencing reads were used, ambiguous bases (N, R, Y, etc.) were discarded. Next, RNAsnp (Sabarinathan et al. 2013) and SNPfold (Halvorsen et al. 2010; Corley et al. 2015) were used to determine which of these SNPs may be riboSNitches (Figure 3, Step 5). These two programs work by generating secondary structure ensembles for the reference and each SNP sequence and quantitatively comparing their partition functions via a correlation coefficient (see Methods section *“Application of RNAsnp and SNPfold to SARS-CoV-2”*) (Halvorsen et al. 2010; Sabarinathan et al. 2013; Corley et al. 2015). Low correlation coefficients between the reference and SNP ensembles correspond to potential riboSNitches.

PseudoSNitches were identified in the next step (Figure 3, Step 6). Briefly, reference and SNP-containing pseudoknots were predicted by SPOT-RNA and ProbKnot (Bellaousov and Mathews 2010); these two programs were chosen because they have similar PPV (Table S6) for test pseudoSNitches (Table S1). The predicted reference and alternative SNP-containing pseudoknots were then compared using our custom CountPseuku program to identify pseudoSNitches (See Supplemental Text Sections 4-5).

Some SNPs that change the structure of the RNA (riboSNitches/pseudoSNitches) also change the amino acid being encoded. In these cases, it is ambiguous whether the selective pressure is acting on RNA structure or protein structure/function. We thus filtered out all missense non-conservative and nonsense mutations in protein coding regions, leaving behind only missense conservative mutations, silent mutations, and mutations in non-coding regions. Although we did not pursue it herein, the filtered-out data could be mined for SNPs that might serve dual roles in changing amino acid identity *and* RNA structure.

### Analysis of the Frameshifting Element (FSE) in SARS-CoV-2 Genome

Programmed (–1) ribosomal frameshifting (PRF) by an RNA pseudoknot-containing FSE is essential to the lifecycle of many viruses (Jacobs and Dinman 2004; Plant et al. 2005; Sun et al. 2021). Indeed, the replication of SARS-CoV-2 can be impeded by PRF-inhibitor small molecules such as merafloxacin (Dinman et al. 1998; Brakier-Gingras et al. 2012; Sun et al. 2021). The three-dimensional structure of the SARS-CoV-2 FSE was recently reported by RNA chemical probing experiments (Manfredonia et al. 2020; Huston et al. 2021), molecular dynamics simulations (Schlick et al. 2020), and crystallography (Zhang et al. 2021). We begin here by considering the canonical conformation of the FSE that we refer to as the “S2 (stem 2) conformation”, which contains the three stems S1, S2 (the pseudoknot stem), and S3 (Figure 4A, B) (Schlick et al. 2021; Lan et al. 2022). Below we consider the alternative “S2′ conformation” that shares the same S1 and S3 helices but swaps pseudoknot stem S2 with the alternate pseudoknot stem S2′ (Figure 4C) while maintaining the majority of one strand of the pseudoknot stem between both conformations as S2_5′_ (≈S2′_3′_).

**Figure 4:**
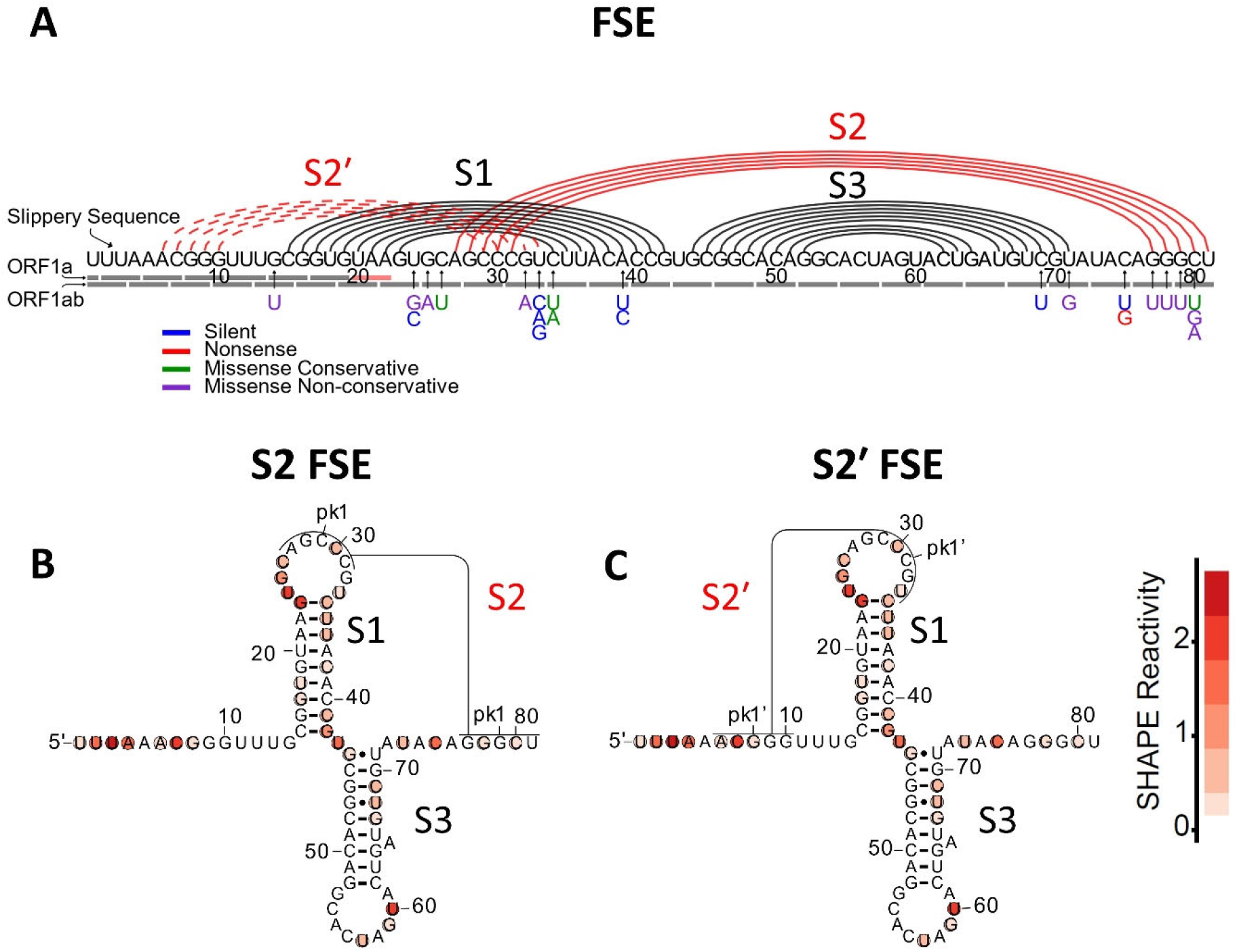
Two structures of the FSE with in vivo chemical probing data. **A)** The FSE can adopt two mutually exclusive conformations. The “S2 FSE” conformation is comprised of three stems: S1 and S3, and the non-nested pseudoknot stem S2, all shown on a circle plot. The alternative “S2′ FSE” conformation is comprised of three stems: the same S1and S3, and a new non-nested pseudoknot stem S2′. The reference sequence is provided along the x-axis, with SNPs shown below. Predicted secondary structures are shown as semicircles, with black representing S1 and S3, and red representing the pseudoknot stem S2 or S2′. The color of each SNP indicates the type of amino acid mutation that occurs when that SNP is present (see legend). **B)** S2 and **C)** S2′ conformations of the FSE with *in vivo* SHAPE data (Huston et al. 2021) superimposed generated using R2easyR (Jacob P. Sieg, Peter C. Forstmeier) and R2R (Weinberg and Breaker 2011). Structures in B and C are from Schlick et al,(Schlick et al. 2021) where the “S2′ FSE” conformation is called “Dual 3_3 FSE”.

We applied the FRID pipeline to the canonical FSE-containing portion of the genome. Initially, we examined the 67 nt stretch of the genome from positions 13476 to 13542, which corresponds to nucleotides 15 to 81 in Figure 4. As mentioned above, RNABOB (Figure 3, Step 1) did not find any motifs. When we applied ScanFold (Figure 3, Step 2) to the genome with the optimal z-score threshold of -2.53σ, the FSE was in fact not identified. This false negative may be due to the FSE adopting multiple stable conformations (Schlick et al. 2020, 2021; Lan et al. 2022), which might lead to a less negative z-score by increasing the denominator (Eq. S1). We then tested if the FSE would score as a true positive upon lowering the stringency of the z-score threshold. Indeed, upon slightly lowering it, to -2.29σ, the FSE was detected. Gratifyingly, SPOT-RNA predicted all five of the accepted FSE pseudoknot base pairs in the S2 conformation (Figure 4A, B) (Schlick et al. 2021; Lan et al. 2022).

Next, we analyzed the FSE in almost 7,000 different SARS-CoV-2 isolates (provided in FRID_viral_isolate_Accessions.xlsx) using BLASTn (Figure 3, Step 4), which identified 23 SNPs at 15 different positions, including some within the canonical and the pseudoknot stems (Figure 4A). As summarized in Table 1, many of these SNPs occurred in unpaired portions of the L (loop), J (joining), and upstream regions, suggesting that these SNPs are not pseudoSNitches because they are common to the S2 and S2′ conformations (see Table 1, SNPs at positions 14, 24, 25, 26, and 75). Other SNPs involved C>U transitions that resulted in GC to G•U changes, which as described above tend to function equally well as base pairs in pseudoknots (see Table 1, SNPs at positions 34, 69, and 80). Some SNPs interfered with pairings in the S1 or S3 helices without shifting the S2 to S2′ equilibrium and were considered pseudoSNitches (see Table 1, SNPs at positions 34, 39, and 71). Five transversion SNPs occurred uniquely in S2_3′_, which is peculiar to the S2 conformation. These SNPs may thus result in enhanced population of the S2′ conformation via weakening of S2 (see Table 1, SNPs at positions 77-80). Similarly, four transversion SNPs occurred uniquely in S2′_3′,_ which is peculiar to the S2′ conformation. This set of SNPs may thus result in an enhanced population of the S2 conformation via weakening of S2′ (Table 1, SNPs at positions 32 and 33). We refer to the changes at 32, 33, and 77-80 as “pseudoknotswitches” because they switch the equilibrium between the two mutually exclusive pseudoknots, not unlike how a riboswitch switches between two mutually exclusive hairpins such as the terminator and antiterminator (Babitzke et al. 2019). Recent computational and experimental data (Schlick et al. 2020, 2021) support the importance of the S2′ pseudoknot conformation of the FSE. These experimental data showed that FSE constructs with greater sequence context than those in the S2 conformation have a frameshifting rate that is nearly double that of the S2 conformation and is closer to that observed in viral infections (Lan et al. 2022).

**Table 1:** SNPs occurring in the FSE of SARS-CoV-2

| Position<sup>a</sup> | Reference | ALT | PseudoSNitch | Mutation Type | Favors S2 or S2' <sup>b</sup> |
| --- | --- | --- | --- | --- | --- |
| 14 (UP) | G | U | No | Missense Non-conservative | Neither |
| 24 (L1)<sup>c</sup> | U | G | No | Missense Non-conservative | Neither |
| 24 (L1) | U | C | No | Silent | Neither |
| 25 (L1) | G | A | No | Missense Non-conservative | Neither |
| 26 (L1) | C | U | No | Missense Conservative | Neither |
| 32<sup>d</sup> (L1) | G | A | PKS<sup>e</sup> | Missense Non-conservative | S2 |
| 33<sup>d</sup> (L1) | U | C | PKS | Silent | S2 |
| 33 (L1) | U | A | PKS | Silent | S2 |
| 33 (L1) | U | G | PKS | Silent | S2 |
| 34 (S1<sub>3'</sub>)<sup>f</sup> | C | U | No (forms G•U) | Missense Conservative | Neither |
| 34 (S1<sub>3'</sub>) | C | A | PSN<sup>g</sup> | Missense Conservative | Neither |
| 39 (S1<sub>3'</sub>) | A | U | PSN | Silent | Neither |
| 39 (S1<sub>3'</sub>) | A | C | PSN | Silent | Neither |
| 69 (S3<sub>3'</sub>) | C | U | No (forms G•U) | Silent | Neither |
| 71 (S3<sub>3'</sub>) | U | G | PSN | Missense Non-conservative | Neither |
| 75 (J3/2)<sup>h</sup> | C | U | No | Silent | Neither |
| 75 (J3/2) | C | G | No | Nonsense | Neither |
| 77 (S2<sub>3'</sub>) | G | U | PKS | Missense Non-conservative | S2' |
| 78 (S2<sub>3'</sub>) | G | U | PKS | Missense Non-conservative | S2' |
| 79 (S2<sub>3'</sub>) | G | U | PKS | Missense Non-conservative | S2' |
| 80 (S2<sub>3'</sub>) | C | U | No (forms G•U) | Missense Conservative | Neither |
| 80 (S2<sub>3'</sub>) | C | G | PKS | Missense Non-conservative | S2' |
| 80 (S2<sub>3'</sub>) | C | A | PKS | Missense Non-conservative | S2' |

<sup>a</sup>Note that the “Position” column is defined for the S2 conformation. <sup>b</sup>Favors the S2 or S2' conformation denoted in Figure 4A and B, respectively. <sup>c</sup>LX denotes the loop region of Stem X (SX). <sup>d</sup>Positions 32 and 33 are contained within S2'<sub>3'</sub>. <sup>e</sup>PKS = Pseudoknotswitch between the S2 and S2' conformations. <sup>f</sup>SX<sub>3'</sub> means the 3' segment of SX. <sup>g</sup>PSN = PseudoSNitch. <sup>h</sup>JX/Y denotes the joining region between SX and SY, lettering 5' to 3'.

It is notable that there were no mutations to the CCC stretch (positions 29-31), which is shared between S2_5′_ and S2′_3′_, (Figure 4A) possibly indicating that both pseudoknot conformations are essential to the viability of the virus. Moreover, *in vivo* SHAPE chemical probing data show strong protection of this CCC stretch (Huston et al. 2021), consistent with its persistent participation in one or the other pseudoknot pairing. Interestingly, protection of S2_3′_ and S2′_5′_ nucleotides, present uniquely in the S2 and S2′ conformations respectively, was only weak, consistent with just partial occupancy and thus their formation being mutually exclusive (Figure 4 B, C) (Huston et al. 2021). Given its conservation, targeting of the FSE pseudoknot by antisense oligonucleotides (ASO) or small molecule therapeutics may be worth considering (Bennett and Swayze 2010; Khvorova 2022). Overall, our data support a functional role of the S2′ conformation of the FSE in frameshifting regulation, as recently proposed (Schlick et al. 2020, 2021), potentially providing an additional target for therapeutics. ASOs could be designed to interfere with either the S2 conformation, S2′ conformation, or both conformations simultaneously.

### Analysis of the Putative Functional RNA Elements in the SARS-CoV-2 Genome

After benchmarking the FRID pipeline on a known pseudoknot containing element, we searched for novel functional RNA elements throughout the entire SARS-CoV-2 genome. As mentioned, ScanFold identified 60 structured regions with a z-score ≤ -2.53σ, 22 of which were predicted to contain a pseudoknot by SPOT-RNA (Figure 2, Table S3). These 22 pseudoknot-containing RNA candidates were then input into the RSC module (Figure 3) to test their conservation using nearly 6,400 viral isolates (provided in FRID_viral_isolate_Accessions.xlsx). All 22 candidate pseudoknot containing RNAs showed levels of conservation equal to or higher than both the nested and pseudoknot base pairs of the S2 conformation of the FSE (see Methods section “*Benchmarking the FRID pipeline on the FSE*”). Eight candidate pseudoknot-containing RNAs contained more nonsense or missense non-conservative mutations than the FSE and were not examined further (Figure 2 and Table S3), which left 14 conserved putatively functional pseudoknots (PFPs) (Table 2). These candidate functional RNAs are enriched in the RdRp and S protein regions of the genome (Figure 2), which is encouraging as these are essential genes in the replication of SARS-CoV-2, making them ideal therapeutic targets. One notable feature of the PFPs is the large number of pseudoSNitches as compared to riboSNitches (Table 2). Not only do SNPs in pseudoknot-forming strands affect pseudoknot formation, SNPs in some of the non-pseudoknot-forming strands also affect pseudoknot formation. It may be the case that pseudoknots are generally more sensitive to SNPs than standard helices.

**Table 2:**
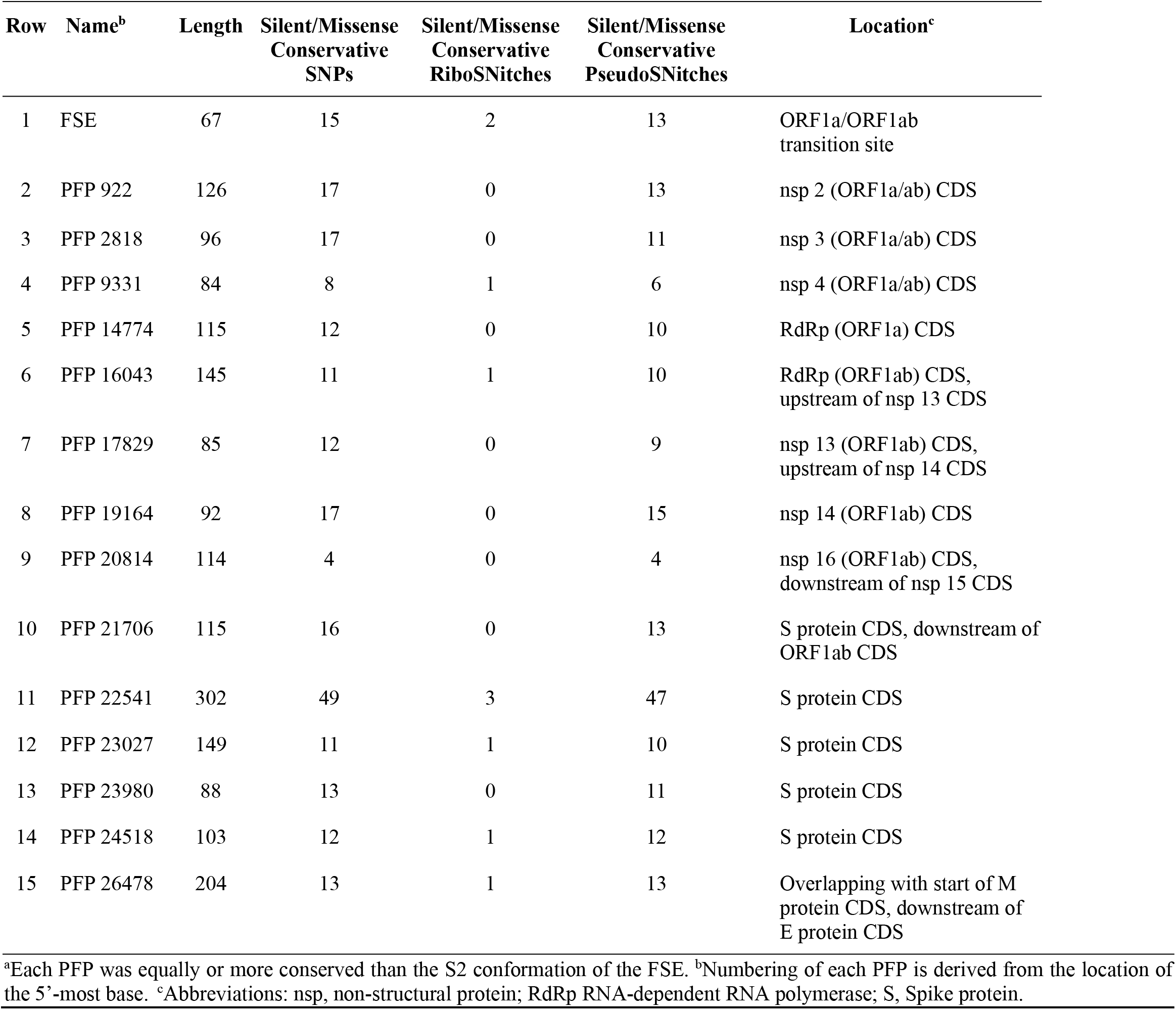
Levels of conservation of the most conserved Putative Functional Pseudokotted elements (PFPs).^a^

One PFP of particular interest is PFP 9331, which is located in nsp 4, a membrane-spanning protein implicated in recruitment of the viral replication-transcription complex (RTC) to modified endoplasmic reticulum membranes (Paysan-Lafosse et al. 2023). The PFP 9331 element folds with a core secondary structure comprised of two putative pseudoknot stems denoted ‘S1’ and ‘S2’ and three stems denoted ‘S3’, ‘S4’, and ‘S5’ (Figure 5A). According to Eq. S3 and the data in Table 2, these stems show lower normalized rates of SNPs, riboSNitches, and pseudoSNitches than the stems of the FSE, indicating that PFP 9331 is a highly conserved region. Moreover, some SNPs potentially strengthen the existing structure by adding canonical base pairs to stems such as the C54G SNP. Transitions like A37G and C61U in S4 and S3, respectively, are not disruptive as they can form G•U wobble pairs, as discussed above. Most of the other SNPs are in single-stranded regions suggesting that they are non-disruptive, with the S2 pseudoknot devoid of SNPs and the S1 pseudoknot having just one disruptive C>G mutation.

**Figure 5:**
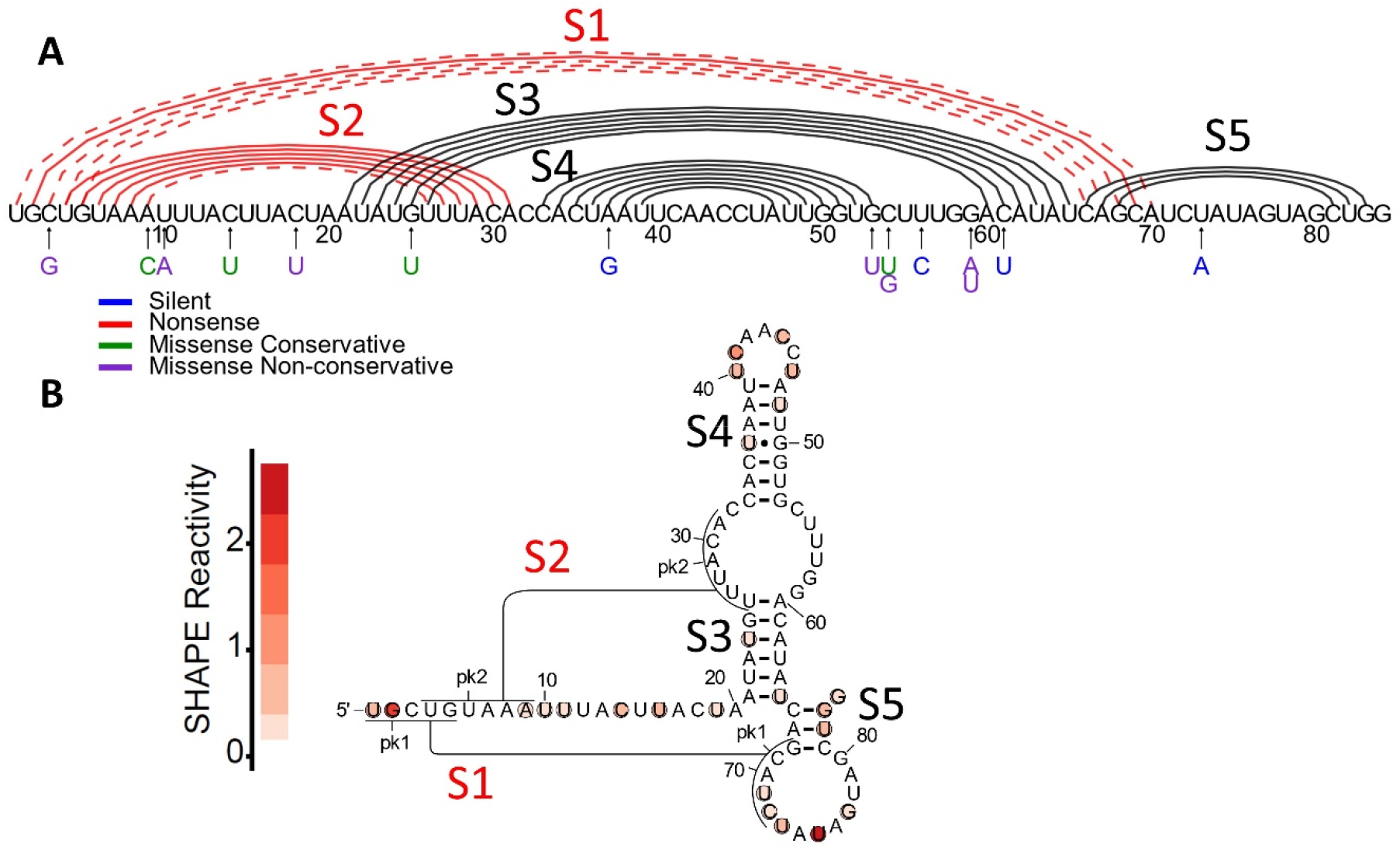
PFP 9331 structure and in vivo chemical probing data. **A)** PFP 9331 shown with the reference sequence across the x-axis. The secondary structure is represented by black semicircles and the pseudoknot by red semicircles. Additional potential pseudoknot pairs in S1 are represented by red dashed semicircles. The color of each SNP shown below the x-axis indicates the type of amino acid mutation that occurs when that SNP is present (see legend). **B)** The FRID-predicted structure of PFP 9331 superimposed with *in vivo* SHAPE chemical probing data (Huston et al. 2021) generated using R2easyR (Jacob P. Sieg, Peter C. Forstmeier) and R2R (Weinberg and Breaker 2011).

There are two putative pseudoknots in PFP 9331. The 5′-most pseudoknot stem, S1, was predicted to contain only a single base pair but the 3′-most pseudoknot, S2 has five or six canonical base pairs, albeit only a GC pair. The unique combination of the high PPV and low sensitivity of SPOT-RNA (Table S6) means that it often does not predict the entire pseudoknot, but when it does predict a pseudoknot base pair it is usually correct. And this means that there may be additional undetected base pairs in the predicted pseudoknots, as we see in most of the PFPs. While pseudoknot S1 may seem very weak, this single base pair is flanked by four additional possible members of the stem, three on one side and one on the other, shown via dashed red lines in Figure 5A. These additional pseudoknot base pairs are mutually exclusive with S5 and with two of the five to six base pairs in S2. One possibility is that S2 and S5 break when S1 forms, meaning that PFP 9331 could be a pseudoknotswitch.

To further evaluate the FRID-predicted structure of PFP 9331, we overlaid it with available *in vivo* SHAPE chemical probing data (Figure 5B) (Huston et al. 2021). Not only are the stems generally unreactive and the loops reactive, which supports the secondary structure, but the S2 pseudoknot shows no reactivity and no SNPs, suggesting that it may form much of the time. The S1 pseudoknot, on the other hand, shows partial reactivity and a stem-breaking C>G mutation, which makes its formation less certain.

The lack of destructive SNPs, high structural conservation, and overlap with SHAPE chemical probing data provide strong evidence that the PFP 9331 RNA element likely folds *in vivo*. The remaining 13 PFPs are described in the Supplemental Information (see Supplemental Text Section 6 and Figures S4-S16). Overall, there are three classes of the 14 PFPs. The first class contains high confidence PFPs, in which there are clear stable pseudoknots, transitions to G•U wobbles that maintain pseudoknot base pairing, and consistency with available SHAPE probing data; these include the following 6 sequences: PFP 922, PFP 2818, PFP 9331, PFP 14774, PFP 17829, and PFP 26478. The second class contains moderate confidence PFPs, in which at least one of these three criteria is lacking; these include the following 2 sequences: PFP 16043 and PFP 23027. The third class contains lower confidence PFPs, in which at least two of these criteria are lacking; these include the following 6 sequences: PFP 19164, PFP 20814, PFP 21706, PFP 22541, PFP 23980, and PFP 24518.

As stated above, FRID identified 14 conserved pseudoknotted regions that are as well conserved or better than the well-known pseudoknot in the FSE. Using the 3 criteria provided above, the structures of approximately half of these PFPs were described with high confidence and the other half with low confidence. However, regardless of their agreement with the 3 criteria we provide, all 14 of these PFPs are conserved, highly structured RNAs.

## Conclusions

The FRID pipeline offers an agnostic computational approach for finding novel candidate pseudoknot containing functional RNAs. It evaluates RNA primary structure, secondary structure, and pseudoknot structure conservation. We developed the FRID pipeline and applied it to SARS-CoV-2, where we identified several novel putative pseudoknot containing regions that are potential therapeutic targets for disruption via ASOs or small molecules therapies already validated for disruption of functional RNAs and RNA-protein interactions (Bennett and Swayze 2010; Hargrove 2020; Khvorova 2022; Zhu et al. 2022).

Excitingly, the structures of some of the PFPs described here appear to agree well with available *in vivo* structure probing data and demonstrate RNA structure conservation in the face of SNPs. Experiments leveraging high throughput screening techniques such as massively parallel oligonucleotide synthesis, *in vitro* selection, and chemical probing followed by Next-Generation Sequencing (NGS) could allow for the probing of a multiplicity of sequence variants. Depending on the throughput of the assay, the z-score threshold could then be relaxed to generate more candidates, albeit at the cost of more false positives.

The putative functional RNAs identified herein could play important roles in the regulation of transcription, sgRNA synthesis, or translation, suggesting that their unfolding could inhibit viral replication. Due to their extensive structures, these RNAs are likely to be vulnerable to ASO or small molecule therapy, providing new avenues for development of RNA-targeted antiviral therapeutics.

## Methods

### PK Module

#### Application of RNABOB to SARS-CoV-2

Descriptor files (.des) were written for the structural motifs that were searched for with RNABOB (Eddy 2012). These files serve as rules when searching for motifs in a genome or transcriptome. The descriptor files can be written to allow additional base pairs, bulges, and variable loop lengths in the search motif. For example, in some of the descriptors used herein, some stems were allowed to be up to 4 bp longer and include up to two mismatches. Descriptors for the HDV, hammerhead, and twister ribozymes, and the guanine, adenine, and deoxyguanosine riboswitches were used (see Supplemental Descriptors for the exact descriptors). No RNABOB results were found in SARS-CoV-2 for the motifs that were searched.

#### Application of ScanFold to SARS-CoV-2

ScanFold has two separate programs that work together, ScanFold-Scan and ScanFold-Fold (Andrews et al. 2018), which were piped together in the FRID pipeline. ScanFold-Scan constructs a file with the z-score data for all predicted *i-j* pairs from the inputted fasta file using the optimized parameters of a sliding window of 120 nt with a step size of 10 nt (Andrews et al. 2018). The z-score was obtained by subtracting the reference RNA free energy of folding and the normalized average stability of 50 randomly shuffled sequences that retain dinucleotide content and normalizing by its standard deviation (Eq. S1). These results were piped directly into ScanFold-Fold which returns the final predicted base pairs of the entire input sequence after it combines the overlapping 120 nt sliding window frames. We then sorted these base pairs into local structured regions by combining any helices that were within 10 nt of each according to our optimization. This distance was optimized using series of known controls from Rfam (Kalvari et al. 2021) (Table S1). The positions of the beginning and end of each of these structures were also adjusted according to an optimization for ScanFold biases. The adjusted beginning and end positions were used in all further analyses.

#### Application of SPOT-RNA & CountPseuku to SARS-CoV-2

While SPOT-RNA (Singh et al. 2019) utilizes parallelization in its computation, it is still exceedingly computationally expensive. Therefore, we input only candidate structures found by ScanFold, instead of applying it to the entire genome. The predicted structure generated by SPOT-RNA was input into CountPseuku (See Supplemental Text Section 5) to determine if there were pseudoknots in the structure. According to the standard CountPseuku procedure, stems were constructed, and the number of pseudoknot stems were determined.

### RSC Module

#### Application of BLASTn to SARS-CoV-2

BLASTn (Morgulis et al. 2008) was used to identify and index naturally occurring SNPs. A large library of SARS-CoV-2 complete genome isolates was downloaded from NCBI and GSAID (n=6362 for all PFPs, n=6975 for the FSE, see FRID_viral_isolate_Accessions.xlsx). No strain specificity was used in collecting these isolates. Each isolate was automatically compared to the reference sequence using BLASTn, and the SNPs were recorded in a text file.

#### Application of RNAsnp and SNPfold to SARS-CoV-2

RNAsnp and SNPfold were used to identify riboSNitches (Halvorsen et al. 2010; Sabarinathan et al. 2013; Corley et al. 2015). In these programs, secondary structure ensembles were generated via a partition function for the reference and each SNP sequence for each 120 nt window. These two partition functions were compared, and the SNP was considered a riboSNitch if its ensemble was significantly different than the reference ensemble or if its population distribution differed from the reference ensemble according to the criteria provided below. The significance of this difference was measured by comparing the two partition functions and calculating a correlation coefficient. Two different riboSNitch-detecting programs, RNAsnp and SNPfold, were used as they have been shown to perform with similar rates of success as binary classifiers by ROC analyses, and the computational cost of employing both was low (Corley et al. 2015). In cases where only one of the two programs predicted a riboSNitch, the pipeline still considered it a riboSNitch and analysis continued.

Each SNP that was input into RNAsnp was folded with up to 60 nt of the native sequence flanking either side of the SNP whenever possible. If the SNP appeared within 60 nt of the end of a candidate region, owing to being near the end of the element, determined by ScanFold, the folding window was truncated to the edge of the candidate region. RNAsnp outputs two values, the local Pearson correlation coefficient (r_min_) and the local Euclidean distance (d_max_). If either the r_min_ or d_max_ p-values were ≤ 0.1, then the SNP was considered a riboSNitch.

Likewise, each SNP input into SNPfold was folded with up to 60 nt of the native sequence flanking either side of the SNP. The key output of SNPfold used to determine if a SNP was a riboSNitch was the Pearson correlation coefficient of the base pairing probability (BP_prob_). If the BP_prob_ was ≤ 0.8, then the SNP was considered a riboSNitch.

#### Application of ProbKnot, SPOT-RNA and CountPseuku to SARS-CoV-2

Two sequence windows were constructed for every SNP: one was the alternative (SNP) sequence with up to 60 nt flanking the SNP on either side, and the second was the reference sequence with the same sequence context. Both sequences were input into ProbKnot and the predicted structures of the alternative (SNP) and the reference sequences were compared with CountPseuku, which determined the number of pseudoknot base pairs and their locations in the two structures. Any difference in the number of pseudoknot base pairs or their locations was considered a pseudoSNitch. The same procedure was repeated with SPOT-RNA instead of ProbKnot as the structure prediction program. If either program called a pseudoSNitch, the SNP was considered a pseudoSNitch.

#### Benchmarking the FRID pipeline on the FSE

Functional RNAs experience selective pressures to maintain their fold. As such, they are unlikely to experience high levels of SNPs, riboSNitches, or pseudoSNitches (Zúñiga et al. 2009; Wu et al. 2015). We calculated length-normalized rates of SNPs, riboSNitches, and pseudoSNitches for each candidate functional RNA element and compared these values to those of the FSE (Table 2), where the “length-normalized rates of SNPs” is the number of SNPs in a structure divided by the number of isolates searched, divided by sequence length (Eq. S3). Length-normalized rates of riboSNitches and pseudoSNitches were calculated in the same fashion. Because of its known function in viral replication, the FSE was analyzed using a slightly larger library of viral isolates than the rest of the PFPs (see FRID_viral_isolate_Accessions.xlsx), and the rates of conservation were normalized to library size.

#### Folding of predicted structures using RNAStructure

The mean free energy (MFE) of each structure was determined and read out using the RNAstructure programs Fold and efn2 (Reuter and Mathews 2010). We also provide the ΔΔG in FRID_SNPs_archive.csv, which was calculated by taking the difference of the reference and SNP structures.

#### Implementation of the FRID pipeline on supercomputing clusters

The RSC module of the FRID pipeline was implemented on the Roar supercomputing clusters at the Pennsylvania State University. Due to the differing software requirements of component programs, a slightly altered architecture was implemented on Roar. The main body of the RSC module – including BLASTn, RNAsnp, ProbKnot, SPOT-RNA, CountPseuku, and RNAstructure – were all run from the same script because they all function in Python3. However, SNPfold requires Python2 and so was implemented in a secondary script run in the same batch as the rest of the module. This adjustment required generation of intermediate files with partially complete results to be piped into SNPfold after the rest of the data had been collected. All code for the FRID pipeline is available at https://github.com/forstmeier-pc/FRID.

## Acknowledgments

This work was supported in part by NIH grant 5R35GM127064 and NIH Supplemental Grant 3R35GM127064-03S1.

